# Exploring phosphoregulation of MYO3A using quantitative fluorescence image analysis in COS7 cells

**DOI:** 10.64898/2026.05.05.723000

**Authors:** Vu Hao Minh Nguyen Phan, Omar A. Quintero-Carmona

## Abstract

Myosin 3A (MYO3A) is an unconventional myosin involved in the formation and maintenance of hair-cell stereocilia of the sensory epithelia in the inner ear. The kinase domain has been implicated in phosphoregulation of MYO3A activity through intermolecular autophosphorylation. Previous studies using mass spectrometry identified two potential phosphorylation sites in the motor domain. To investigate the regulatory roles of these sites, we generated glutamic acid point mutations in our mchr-MYO3AΔK construct to mimic phosphorylation and assayed the constructs for their ability to tip-localize and influence filopodial density via transfection into COS7 cells. The phosphomimic constructs were less able to generate filopodia when compared to wildtype constructs. To gain a better understanding of the phosphoregulation of MYO3A, we transfected COS7 cells with mchr-MYO3AΔK in combination with GFP-tagged full-length MYO3A (GFP-MYO3A^FL^), or GFP attached to just the kinase domain of MYO3A (GFP-MYO3A^KIN^). Coexpression of mchr-MYO3AΔK with either construct resulted in decreased mchr-MYO3A levels at the tips of filopodia and fewer filopodia at the edge of the cell, compared to cells expressing mchr-MYO3AΔK alone. This implies that the kinase domain does not require motor activity to contribute to phosphoregulation of MYO3A, and that MYO3A phosphoregulation may be influencing filopodia initiation. Informatic analyses and structural predictions suggest that the two phosphorylation sites in the motor domain inhibit actin/MYO3A interactions. Taken together, these analyses link MYO3A phosphorylation with the regulation of its ability to create actin protrusions such as filopodia and stereocilia.

## Introduction

### Hearing as a biological process

Mammals have the unique ability to sense, amplify, and discern sounds of different frequencies and intensities in the biological process that is hearing (Schwander et al., 2010). This process has much to do with the unique properties of the auditory system, in particular the structures of the inner ear (illustrated in Figure 1). The snail-shaped cochlea lies within the inner ear and is responsible for converting frequencies captured by the outer ear into electrical signals that are then perceive as sound (Casale et al., 2023). Mechanosensory hair cells are essential to this process, and are a component of the organ of Corti--along with additional supporting cells and accessory extracellular structures (Schwander et al., 2010).

**Figure 1:**
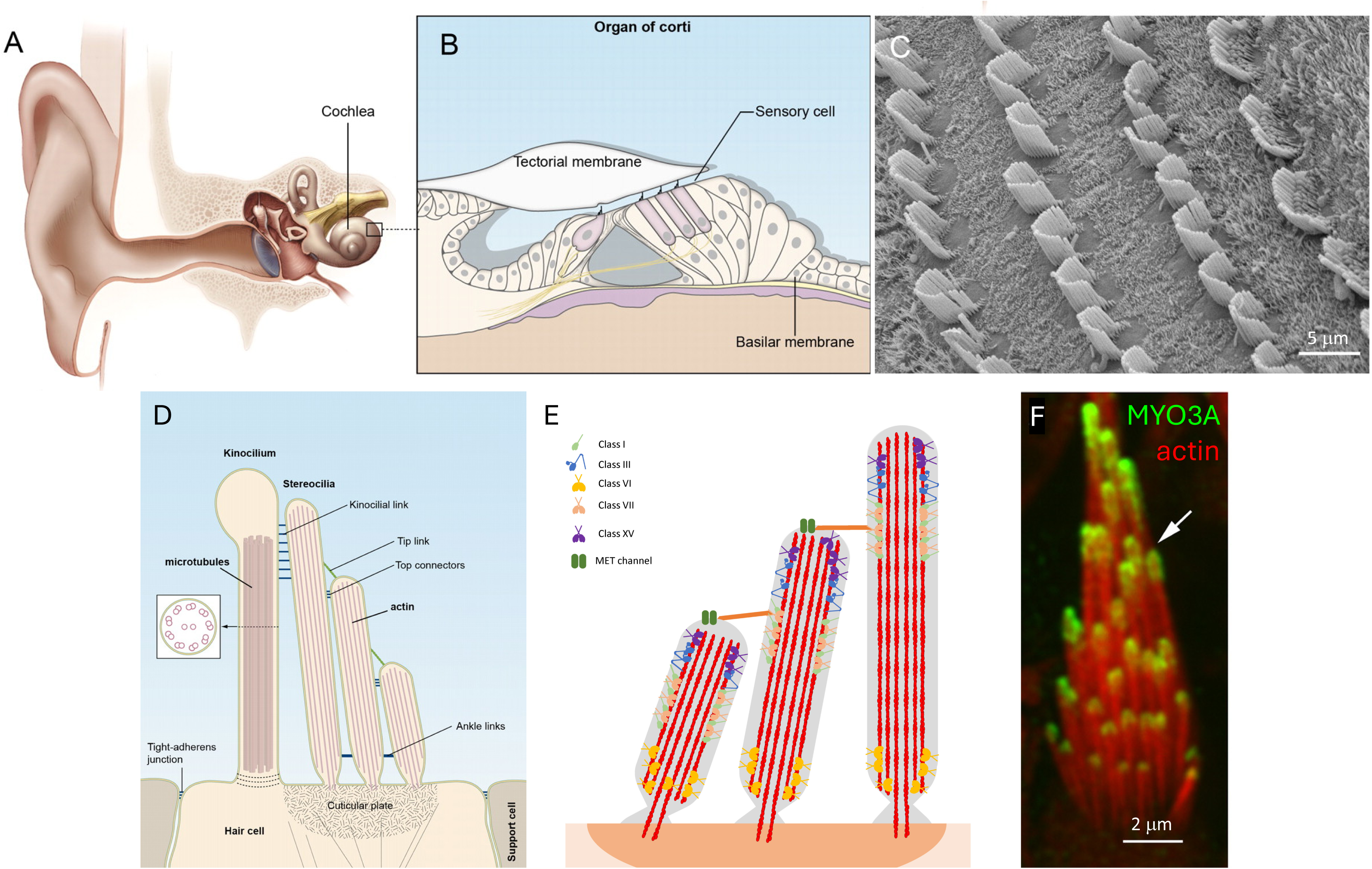
Representations of hearing machinery from the literature. **(A)** Inside the inner ear, the snail shaped cochlea can be found, which is where **(B)** the organ of Corti is located. **(C)** Upon removal of the tectorial membrane, rows of outer hair cells (3 on the left) and inner hair cells (one on the right) can be visualized by scanning electron microscopy**. (D)** When viewed in cross section, the apical hair cell surface is composed of the kinocilium, stereocilia, and cuticular plate. **(E)** Stereocilia are linked together by tip links which control mechanically gated ion channels. Different myosins are distributed to three different regions of the structure, with class III myosins found at the tips. **(F)** Immunostaining reveals MYO3A localized to the tips of stereocilia in to have some immunoreactivity at the tips of stereocilia in the bullfrog inner ear. Panels A-D adapted from (Schwander et al., 2010), and panel E adapted from (Cirilo et al., 2021). Panel F is from (Schneider et al., 2006), copyright 2006, Society for Neuroscience.

The organ of Corti is located within the cochlea and composed of ∼16,000 hair cells that are patterned into one row of inner hair cells (IHCs) and three rows of outer hair cells (OHCs) (Schwander et al., 2010). Each hair cell is composed of a mechanically sensitive organelle known as the hair bundle, which itself is composed of dozens of actin protrusions known as stereocilia (McPherson, 2018). These protrusions are assembled in rows of decreasing height (Figure 1), where the tips of these stereocilia point away from the center of the cochlea in order to maximize the probability of mechanotransduction (Hudspeth & Corey, 1977). OHCs use these stereocilia bundles to form connections with the tectorial membrane that covers the apical surface of the organ of Corti (McPherson, 2018). The cell bodies of the hair cells form tight connections with support cells, which in turn adhere to a structure known as the basilar membrane (McPherson, 2018).

The initiation of hearing occurs when oscillations in air pressure are captured by the outer ear, which then are converted into fluid pressure that travels down the cochlear duct of the inner ear (Schwander et al., 2010). This results in vibrations in the basilar membrane that are transferred to the hair cells located above. These vibrations cause the deflection of the hair bundles leading to the opening of mechanically gated ion channels and the depolarization of hair cells, resulting in neuronal sensation of hearing (Schwander et al., 2010). The proper maintenance of stereocilia is pivotal, especially their height. Similarly to other actin-based protrusions, stereocilia are supported by bundles of uniformly polarized, dynamic actin filaments (Tilney et al., 1992). As hair cells are not replaced if they die, it is vital that the architecture of stereocilia is maintained at a constant length throughout life (Narayanan et al., 2015). To maintain this architecture, the rate of actin polymerization and depolymerization must be tightly coordinated in stereocilia, which is in part, accomplished via expression of cytoskeletal regulatory proteins, including a number of different myosin genes.

### Myosins and hearing

Multiple myosins are involved in the structure and function of stereocilia. As a whole, the myosin superfamily is a large and diverse protein family that is involved in a variety of biological processes including muscle contraction, cell motility, cytokinesis, and hearing (Hartman & Spudich, 2012). Myosin genes are grouped into classes based on phylogenetic analyses of the highly conserved N-terminal motor domain (Cheney et al., 1993). This motor domain converts the energy from ATP hydrolysis into force and motion through a cyclic interaction with actin filaments (Lymn & Taylor, 1971; Ruppel & Spudich, 1996). Additionally, each myosin is characterized by a unique C-terminal tail domain. This tail domain allows the myosins to form interactions that enable them to have individualized cellular functions through association with other proteins or cargos (Akhmanova & Hammer, 2010). With respect to actin-based protrusions from the cell surface, the interactions through the tail domain and the force generation by the motor domain for a subset of specific myosins leads to the formation and maintenance of stereocilia (Nambiar et al., 2010).

At least six myosins implicated in the formation and maintenance of stereocilia, and mutations to these genes have been linked to multiple different forms of deafness (Friedman et al., 2020). Typically, most mutations lead to autosomal recessive non-syndromic hearing loss, abbreviated as DFNB (Duman & Tekin, 2012). The conventional myosins, myosin heavy chain IX (MYH9) and myosin heavy chain XIV (MYH14) are the least understood (Donaudy et al., 2004; Pecci et al., 2018). Unlike other myosins, MYH9 is expressed throughout the stereocilium, appearing in the base as well as within the shaft of stereocilia (Lalwani et al., 2008). Notably, mutations in the motor domain of MYH9 have been linked to nonsyndromic deafness DFNA17 (Kim et al., 2017; Pecci et al., 2008), which could be a result of stereocilia related deformities. Similarly, mutations in MYH14 have been linked to nonsyndromic deafness DFNA4A as well as progressive sensorineural hearing loss (Choi et al., 2011) possibly due to stereocilia damage. The functions of MYH14 and MYH9 in stereocilia and hearing remain active areas of research.

The unconventional myosin, myosin VI (MYO6) is special in the way it moves across actin filaments. Typically, myosins move towards the plus ends of F-actin, but uniquely, MYO6 moves towards the minus ends of F-actin. This phenotype has been attributed to the 53 amino acid insert within the converter domain (Wells et al., 1999). Generation of chimeric myosins through the insertion of the MYO6 converter domain into MYO5 turned MYO5 into a minus end directed motor (Park et al., 2007). In stereocilia, MYO6 is localizes to the base of stereocilia (Avraham et al., 1997). Even in overexpression studies, MYO6 stays consistent in its position at the base of stereocilia (Belyantseva et al., 2005). Hearing loss DFNB36 and DFNB22 have been associated with pathogenic variants of MYO6 (Melchionda et al., 2001). This could be attributed to the fusing of adjacent stereocilia observed in the absence of functional MYO6 (Friedman et al., 2020). The localization of MYO6 and the collapse of stereocilia structure in the absence of MYO6 suggests that the motor may play a role in anchoring the actin-rich cytoskeleton structure, cuticular plate, which would prevent nearby stereocilia from fusing (Friedman et al., 2020).

Myosin VIIA (MYO7A) is found at the upper end of tip links between stereocilia (Grati & Kachar, 2011). These tip links are extracellular filaments that are responsible for “pulling” open the mechanotransduction channels during the vibrations that induce deflection of stereocilia bundles (Grati & Kachar, 2011). Given this, MYO7A has been hypothesized to modulate the mechanoelectrical transduction (MET) in hair cells (Friedman et al., 2020). Knock-out or mutation studies regarding MYO7A or its binding partners have supported this idea due to the reported loss of tension in the MET machinery (Corns et al., 2018; Grillet et al., 2009; Kros et al., 2002). Interestingly, like MYO6, MYO7A can also be found at the base of stereocilia, where it is implicated to play a role in the complex that forms ankle links (Michalski et al., 2007). These links connect the stereocilia to the base of the hair cell but have been noted to disappear by the time of hearing onset in rodents (Goodyear et al., 2005). It is thought that these ankle links hold the hair bundle together as the stereocilia rootlets are being formed so that the hair bundle is properly formed (Kitajiri et al., 2010). Like other myosins, mutations in MYO7A have been implicated in nonsyndromic hearing loss, DFNA11 (X.-Z. Liu et al., 1997), recessively inherited deafness, DFNB2 (Gibson et al., 1995), and most notably, it is the only myosin implicated in Usher syndrome (X. Liu et al., 1997; Weil et al., 1996, 1997). Whirlin, a scaffolding protein and molecular cargo of Myosin XV, is also implicated in Usher syndrome (Ebermann et al., 2007).

Myosin XV (MYO15), and by extension whirlin, can be found at the tips of stereocilia (Belyantseva et al., 2003). The localization of MYO15 at the tips of stereocilia has been implicated to play a very pivotal role in maintaining the lengths of these protrusions to preserve the staircase arrangement of stereocilia bundles (Anderson et al., 2000; Belyantseva et al., 2003). As such, pathogenic mutations in MYO15 have been associated with deafness DFNB3 (Probst et al., 1998; Wakabayashi et al., 1998, p. 2; Wang et al., 1998). However, MYO15 does not seem to play a role in the emergence, initiation, or increase in thickness of stereocilia, playing exclusively a maintenance role within the structure (Belyantseva et al., 2005). This is presumably due to the motor’s role in transporting whirlin, with whirlin’s C-terminal PDZ domain binding to the MYO15’s C terminal PDZ-ligand (Belyantseva et al., 2005). Importantly, the presence of whirlin is not necessary for MYO15 to localize to the tips of stereocilia, but on its own, the motor is insufficient for stereocilia elongation (Belyantseva et al., 2005). Similarly, without MYO15, whirlin expression is insufficient for the elongation of stereocilia (Mburu et al., 2003). Furthermore, in the outer hair cells, the mechanotransduction complex located at the tips of stereocilia form independently from MYO15 mediated stereocilia elongation (Stepanyan & Frolenkov, 2009).

### Class Myosin III and Hearing

Class III myosins are also heavily involved in the tip-localized mechanotransduction complex in addition to serving a variety of other roles related to stereocilia (Grati et al., 2016). Initially characterized in *Drosophila melanogaster* and named NINAC (Montell & Rubin, 1988), class III myosins are unusual members of the myosin superfamily given their conserved N-terminus kinase domain (Dosé et al., 2007). This kinase domain has been proposed to allow for the regulation of the myosin protein through autophosphorylation (Ng et al., 1996; Quintero et al., 2010, 2013). There are two vertebrate isoforms of class III myosins: MYO3A and MYO3B. Both isoforms localize to the tips of stereocilia, where they are thought to play crucial roles in assembly and maintenance of the protrusion (Dosé et al., 2003). MYO3A and MYO3B contain different functional domains in their tails and are thought to bind to different (and potentially overlapping) subsets of proteins. They have been shown to interact with stereocilia resident proteins like the actin bundling protein espin-1 (ESPN1) (Merritt et al., 2012; Salles et al., 2009) and the adaptor protein MORN4 (Mecklenburg et al., 2015). MYO3A is hypothesized to transport both proteins along stereocilia and is thought to play a pivotal role in regulating stereocilia lengths (Dosé et al., 2003). MYO3A also has an actin-binding domain in its tail, which could allow MYO3A to crosslink actin filaments and promote polymerization-based formation of actin protrusions (Les Erickson et al., 2003). Additionally, MYO3A has been reported to transport Protocadherin 15-CD2, a vital MET protein, to the tips links of stereocilia (Grati et al., 2016). Inactivating mutations in the *MYO3A* gene leading to the loss of function of the protein have been linked to nonsyndromic deafness (DFNB30) (T. Walsh et al., 2002). Mouse lines modeling the DFNB30 mutation display age-dependent hearing loss due to degeneration of hair cell stereocilia (V. L. Walsh et al., 2011).

Immunostaining of MYO3A shows strong labeling of the tips of stereocilia and GFP-MYO3A constructs also show strong tip-localization (Schneider et al., 2006). Fluorescently-tagged MYO3A constructs localize to the tips of other actin-based protrusions such as filopodia (Les Erickson et al., 2003; Schneider et al., 2006) and microvilli (An et al., 2014; Raval et al., 2016) in multiple cultured cell types. As MYO3A contains a kinase domain N-terminal of the rest of the protein (Dosé & Burnside, 2000), the scientific community suspected that MYO3A activity could be regulated via autophosphorylation (Komaba et al., 2003, 2010). Kinase domain deletion (MYO3AΔK) led to enhanced tip-localization of GFP-MYO3AΔK constructs in both hair cell stereocilia (Schneider et al., 2006) and COS7 cell filopodia (Salles et al., 2009). Additionally, introducing a kinase-inactivating mutation into full length GFP-MYO3A also led to enhanced tip localization in hair cell stereocilia and COS7 cell filopodia (Quintero et al., 2010). Ectopic expression of fluorescently-tagged MYO3AΔK constructs in COS7 cells results in an increase in the number of filopodia extending from the edge of the cell (Quintero et al., 2010). The MYO3A kinase domain is capable of intermolecular autophosphorylation *in vitro* (Quintero et al., 2010), and when kinase-functional GFP-MYO3A^FL^ constructs are coexpressed with kinase-deleted mchr-MYO3AΔK constructs, the filopodial initiation phenotype is minimized (Quintero et al., 2010). One current model for MYO3A function imagines that autophosphorylation of the kinase domain happens at the tips of existing actin protrusions and controls the available concentration of MYO3A in that compartment by negatively influencing motor activity via autophosphorylation of the motor domain and the rest of the MYO3A protein (Quintero et al., 2013).

Since the current thinking is that autophosphorylation of MYO3A by its kinase domain tunes down MYO3A activity, characterizing the mechanism by which MYO3A autophosphorylation regulates the motor domain is vital for understanding its role in normal sensory cell physiology and perhaps in the pathology of MYO3A loss-of-function hearing loss. Phosphoproteomic analysis of *in vitro* phosphorylated human MYO3A identified a number of putative phosphorylation sites—at T184 in the kinase domain, at T908 in the motor domain, and at T919 in the motor domain (Quintero et al., 2013). The web resource, Phosphosite Plus, lists 18 additional putative phosphorylation sites spread across the entirety of the protein (Hornbeck et al., 2015). Although the consequences of phosphorylation of the MYO3A motor domain have been studied using *in vitro* steady-state assays and cell-based assays (Komaba et al., 2010; Quintero et al., 2013), the potential biological function of specific phosphorylation of T908 or T919 phosphorylation has yet to be directly investigated. We generated point mutations in mchr-MYO3AΔK construct to mimic phosphorylation, replacing threonine with aspartic acid (mchr-MYO3AΔK^T908D^ and mchr-MYO3AΔK^T919D^), and applied quantitative imaging techniques to elucidate the biological significance of such posttranslational modifications on the actin protrusion-related functions of MYO3A.

## Materials and Methods

### Expression Plasmids

mchr-MYO3AΔK, GFP-MYO3A^FL^ plasmids had been generated previously (Quintero et al., 2010). The phosphomimic mchr-MYO3AΔK^T908D^ and mchr-MYO3AΔK^T919D^ constructs were generated by performing site-directed mutagenesis on the mchr-MYO3AΔK construct. The GFP-MYO3A^KIN^ construct was generated using megaprimer PCR mutagenesis (Attwell et al., 2003) to move the human MYO3A kinase domain (amino acids 1-351) into pEGFP-N1 (Clontech). All expression plasmids were sequence-verified. A schematic of these constructs can be found in Figure 2.

**Figure 2:**
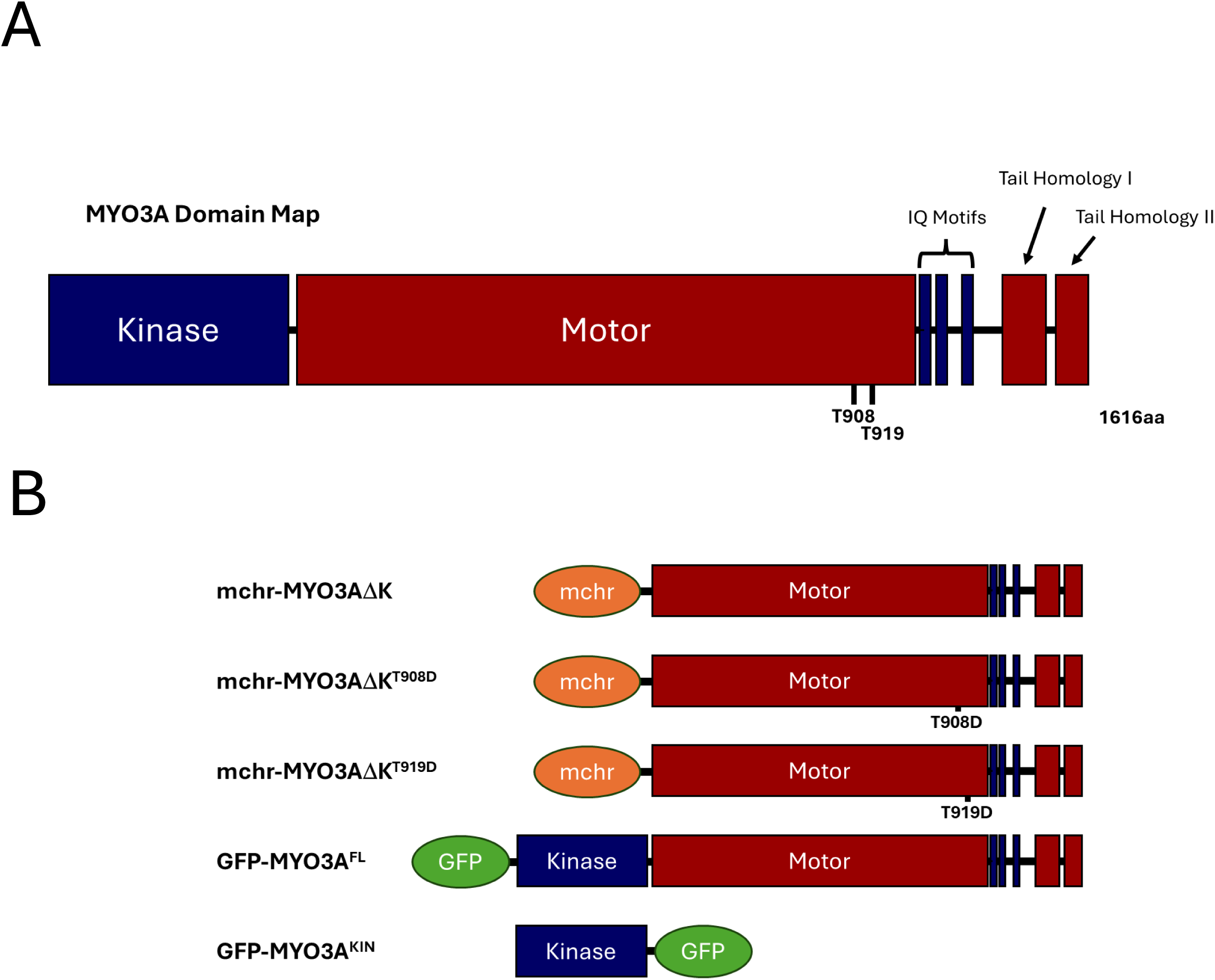
Schematic diagram of MYO3A domains and MYO3A constructs used in this study. **(A)** Human MYO3A is a 1616 amino acid protein consisting of multiple domains in addition to an N-terminal kinase domain followed by a myosin motor domain. The tail domains mediate interactions with other proteins such as F-actin, ESPN1, and MORN4. **(B)** For these studies, human MYO3A sequence was combined with mcherry or GFP sequence to generate specific expression constructs.

### COS7 cell culture and transient transfection

COS7 (Gluzman, 1981) cells were cultured in growth media: DMEM high glucose (Gibco) that was supplemented with 10% Fetal Bovine Serum (Benchmark FBS, GeminiBio), and antibiotics (50U/mL penicillin, 50µg/mL streptomycin, Gibco). Cells were kept in a humidified incubator at 37°C, with a concentration of 5% CO_2_, and passaged using 0.25% trypsin-EDTA (Gibco). In preparation for transfection, cells were plated onto acid-washed, 22mm^2^, #1.5 coverslips at a concentration of ∼30,000 cells/coverslip (one coverslip per well in a six well dish) and then allowed to adhere overnight. Cells were transfected using Lipofectamine 3000, according to manufacturer’s protocol. For each sample well, 0.3µg of plasmid DNA was diluted into 125μL of Opti-MEM media (Invitrogen) without serum or antibiotics and mixed with 3μL of P3000 reagent. In a separate tube, 4μL of L3000 reagent was diluted into 125μL Opti-MEM. The two tubes were then combined, vortexed and incubated at room temperature for 15 minutes prior to dropwise addition to the sample well. Following transfection, the cells were allowed to grow overnight (16-24h) prior to fixation and counterstaining. For co-transfections, a total of 0.6µg of plasmid DNA is used instead.

### Fixed cell sample preparation and imaging

Samples were fixed for 20 minutes in PBS containing 4% paraformaldehyde, permeabilized for 5 mins in PBS containing 0.5% Trition X-100, and counterstained in PBS containing 6.6nM ALEXA647 phalloidin and 10nM DAPI for 30 minutes. The samples were then washed four times for 5 minutes with PBS. Coverslips were mounted onto slides with ProLong Glass Antifade and allowed to dry. Images were obtained using an Olympus IX-83 microscope with a 60x/1.4NA objective, Sutter shutters & filter wheels, Sedat quad filter set (Chroma), and a Hamamatsu ORCA Flash 4v2 camera. The microscopy hardware and image acquisition settings were controlled by Metamorph software. Typical exposures times were 200ms for ALEXA647 phalloidin and 300ms for GFP or mcherry fluorescence channels. Exposures ranged between 2-10ms. Images were collected in all four channels for all treatment groups.

### Tip to Cell Body Ratio and Filopodia Density Quantification and Analysis

FIJI was used for image analysis (Schindelin et al., 2012). The ratio of tip intensity to cell body intensity (TCBR) was calculated by using 4×4 pixel regions-of-interest to determine the intensity of the background (I_b_), the filopodial tip (I_t_), and the cell body (I_cb_). The ratio of the I_t_ to I_cb_ was calculated after subtracting I_b_ from those measurements using the following equation:

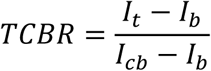

Filopodia edge density was calculated from regions transfected cell periphery not in contact with other cells (free cell edge). Lengths of free cell edge were measured and the number of filopodia along those lengths were counted manually. Filopodia edge density is then calculated as the number of filopodia divided by the length of the measured free cell edge. TCBR and filopodial edge density for each experimental condition were averaged for each day with each day serving as a biological replicate. Statistical significance was determined by ANOVA followed by a Tukey test based on the set of daily averages for each condition. Mean cell brightness was not significantly different between any of the mchr-tagged constructs, nor was mean cell brightness significantly different between any of the GFP-tagged constructs used in this study (Figure 3).

**Figure 3:**
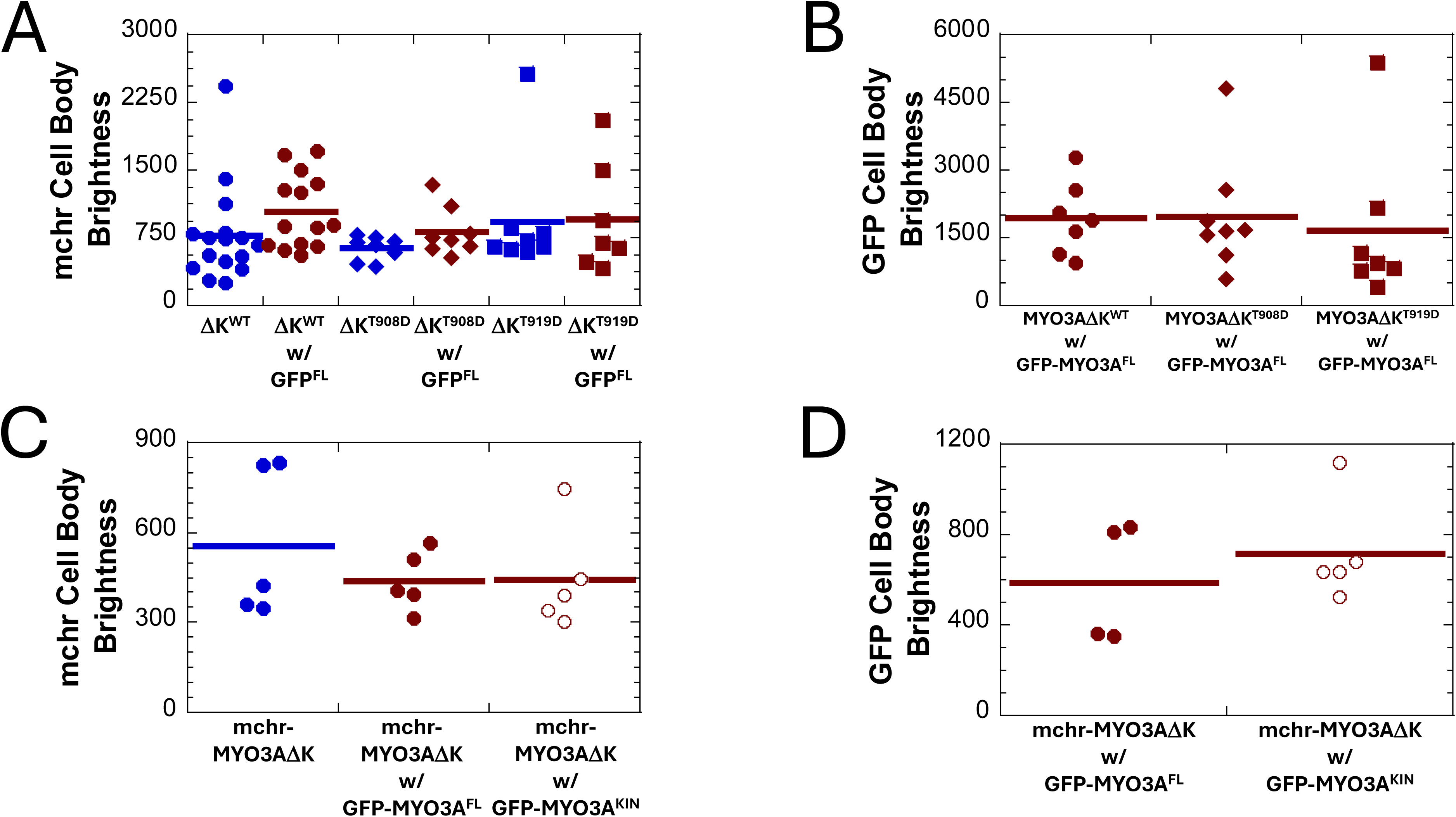
Mean cell body brightness did not vary between expression constructs under the different expression conditions used. This study employed ectopic expression of fluorescently-tagged mchr-MYO3A constructs in COS7 cells. For some experiments, GFP-tagged constructs were also expressed. Mean cell body brightness was measured as part of the tip-to-cell-body-ratio (TCBR) calculation. **(A)** Cell body brightness was not different between the wildtype or phosphomimic constructs whether expressed alone or in combination with GFP-MYO3A^FL^. **(B)** The cell body brightness of the GFP-MYO3A^WT^ did not vary between conditions where it was co expressed with any of the mchr-MYO3AΔK constructs. **(C)** Mean cell body brightness was not different for mchr-MYO3AΔK when expressed alone, in combination with GFP-MYO3A^WT^, or in combination with GFP-MYO3A^KIN^, and **(D)** the mean cell body brightness for the GFP constructs was also not different for those samples. Markers represent biological replicates (independent experiments) and bars represent the mean.

### Phyre Modeling and Protein Sequencing

To predict the tertiary structure of MYO3A, amino acids 316-1616 of the human MYO3A sequence were submitted into the Protein Homology/analogy Recognition Engine (Phyre2.2) web server. Given the sequence, the engine uses homology identification and secondary structure prediction to determine structural homologues by a profile-profile alignment algorithm (Powell et al., 2025). High scoring alignments are then used to construct a proposed tertiary structure of the submitted sequence by comparison to structural databases.

Alternatively, unknown structures can be fit directly to a specific, known structure, which was our approach. As the regions of myosins directly involved in the actin-binding interface were not well characterized in structures generated by X-ray crystallography or cryo-electron microscopy at the time that these studies were initially undertaken (Lorenz & Holmes, 2010), Phyre2 was used to predict the structure of the actin-interface of MYO3A through aligning the sequence of human MYO3A directly to the model of the actomyosin interface generated by Lorenz and Holmes (Lorenz & Holmes, 2010).

### Quantification and statistics

Each condition of each cell-based assay was repeated independently 5 to 8 times (a biological replicate). For TCBR, each biological replicate consisted of at least 10 cells with no more than 5 filopodia assayed per cell. At least 7 cells were measured for each biological replicate for filopodia edge density. Kaleidagraph 4.5 was used to calculate descriptive statistics and carry out the ANOVA with Tukey analysis. Quantification presented in the text are written as mean ± standard error of the mean.

### Selection and preparation of sample images for publication

Sample images chosen for the figure were chosen as representations of the patterns observed in the quantified data. The software packages Metamorph, and FIJI were used to generate the pseudo-colored images included in the figures.

## Results and Discussion

We assayed the phosphomimic constructs for their ability to localize to the tips of filopodia within COS7 cells, which do not express MYO3A endogenously and do not generate many filopodia. Notably, there was no difference in the tip to cell body ratio (TCBR) between cells expressing mchr-MYO3AΔK^T908D^ (3.6±0.3, n=8) compared to mchr-MYO3AΔK^WT^ (4.2±0.3, n=8). Furthermore, cotransfection of the mchr-constructs with GFP-MYO3A^FL^ decreased the TCBR for cells coexpressing mchr-MYO3AΔK^WT^ (2.4±0.4, n=7, p<0.001, Tukey analysis vs WT single-expressor) but not for cells coexpressing mchr-MYO3AΔK^T908D^ (3.1±0.3, n=8, p=0.06, Tukey analysis vs T908D single-expressor). Representative images and sample quantification are shown in Figure 4. However, cells expressing mchr-MYO3AΔK^T919D^ in combination with GFP-MYO3AΔ^FL^ (2.2±0.3, n=8) showed a decrease in TCBR compared to cells expressing mchr-MYO3AΔK^T919D^ alone (3.7±0.3, n=8, p<0.01, Tukey analysis). These data are shown in Figure 5. The pattern that we observed in these data suggests that other factors in addition to phosphorylation at these two sites might be influencing MYO3A localization, such as phosphorylation at additional positions along the MYO3A protein. It is also possible that phosphomimicry does not fully recapitulate the activity of truly phosphorylated MYO3A.

**Figure 4.**
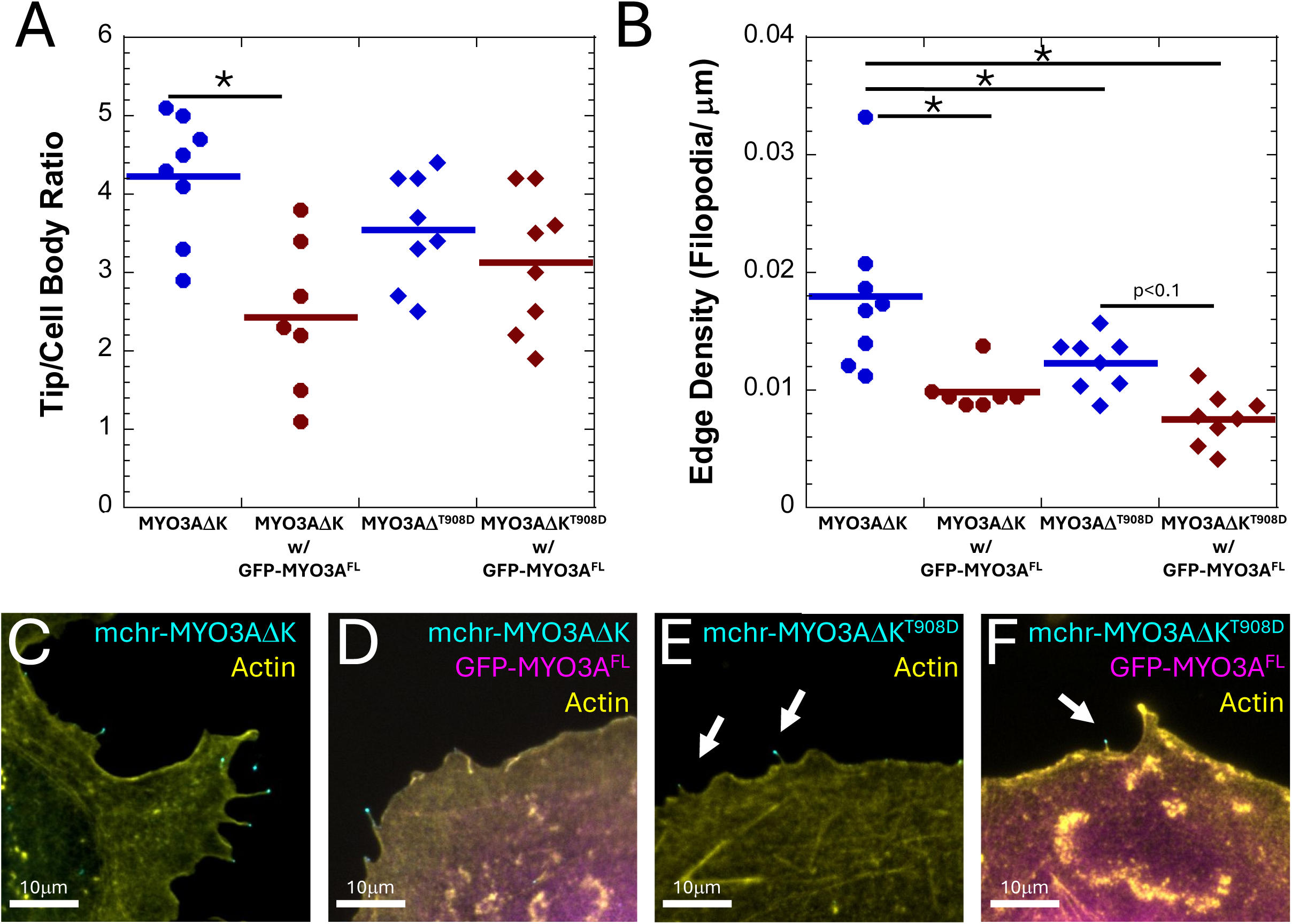
Mimicking phosphorylation of the T908 position in mchr-MYO3AΔK motor results in decrease in filopodial edge density but not tip localization. **(A)** COS7 cells expressing mchr-MYO3AΔK^T908D^ with or without the GFP-MYO3A^FL^ show no difference in tip localization when compared to mchr-MYO3ΔK. Coexpression with GFP-MYO3A^FL^ decreased tip localization for mchr-MYO3AΔK but not mchr-MYO3AΔK^T908D^.**(B)** Filopodial edge density is reduced for cells expressing mchr-MYO3AΔK^T908D^ (with or without GFP-MYO3A^FL^) when compared to cells expressing mchr-MYO3AΔK^WT^. Coexpression of GFP-MYO3A^FL^ with mchr-MYO3AΔK also decreased filopodial edge density. Representative images of transfected cell edges are shown in **C-F**. For all plots, markers represent biological replicates, bars represent the mean, and asterisks represent Tukey analysis with p<0.05.

**Figure 5.**
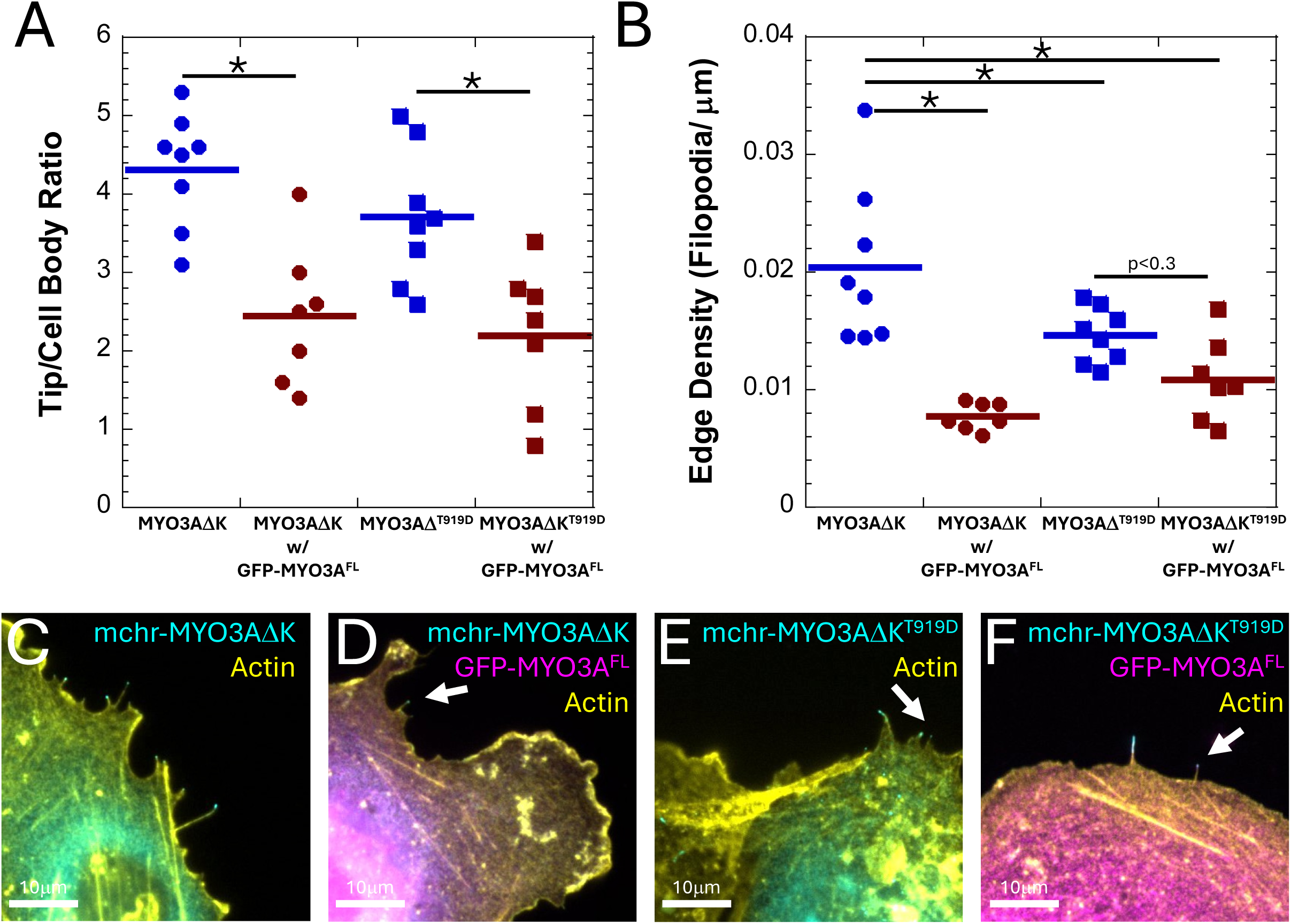
Mimicking phosphorylation of the T919 position in mchr-MYO3AΔK results in decreased filopodial edge density but not tip localization. **(A)** COS7 cells expressing mchr-MYO3AΔK^T919D^ with or without the GFP-MYO3A^FL^ show no difference in tip localization when compared to mchr-MYO3ΔK^WT^. Coexpression with GFP-MYO3A^FL^ decreased tip localization for mchr-MYO3AΔK and mchr-MYO3AΔK^T919D^. **(B)** Filopodial edge density is decreased in cells expressing MYO3AΔK^T919D^ (with or without GFP-MYO3A^FL^) when compared to cells expressing mchr-MYO3AΔK. Coexpression of GFP-MYO3A^FL^ with mchr-MYO3AΔK also decreased filopodial edge density. Representative images of transfected cell edges are shown in **C-E**. For all plots, markers represent biological replicates, bars represent the mean, and asterisks represent Tukey analysis with p<0.05.

We then assayed the individual phosphomimic constructs for their ability to influence filopodial edge density in COS7 cells. There was a significant decrease in the filopodia edge density for cells expressing mchr-MYO3AΔK^T908D^ (0.012±0.0008 filopodia/μm, n=8) compared to cells expressing mchr-MYO3AΔK^WT^ (0.018±0.002 filopodia/μm, n=8, p<0.04, Tukey analysis, Figure 4B). Additionally, coexpression of the mchr-constructs with kinase-functional GFP-MYO3A^FL^ resulted in a statistically significant decrease in filopodial edge density for cells expressing mchr-MYO3AΔK^WT^ (0.010± 0.001 filopodia/μm, n=8, p<0.003, Tukey analysis vs WT single-expressor) but not for cells expressing mchr-MYO3AΔK^T908D^ (0.008±0.001 filopodia/μm, n=8, p=0.1, Tukey analysis vs T908D single-expressor, Figure 4B). The same pattern was observed when comparing cells expressing mchr-MYO3AΔK^WT^ (0.020±0.002 filopodia/μm, n=8), mchr-MYO3AΔK^T919D^ (0.015±0.0008 filopodia/μm, n=8), or those constructs co-expressed with GFP-MYO3A^FL^. Cells expressing mchr-MYO3AΔK^T919D^ had decreased filopodial density compared to cells expressing mchr-MYO3AΔK^WT^(p<0.05, Tukey analysis), and filopodial edge density was decreased when the mchr-MYO3AΔK^WT^ was coexpressed with GFP-MYO3A^FL^(0.008±0.0004 filopodia/μm, n=7, p<0.0001, Tukey analysis vs WT single-expressor). Coexpression of mchr-MYO3AΔK^T919D^ with GFP-MYO3A^WT^ (0.011±0.001 filopodia/μm, n=7) did not significantly decrease filopodial edge density compared to mchr-MYO3AΔK^T919D^ single-expressors (p=0.3, Tukey analysis, Figure 5B). Taken together, these data suggest that phosphoregulation of the MYO3A motor domain influences the ability of MYO3A-expressing cells to support the existence of actin protrusions. Such activity could be the result of MYO3A somehow stabilizing already-formed protrusions, or by stimulating new protrusion initiation events.

If MYO3A autophosphorylation is negatively-regulating the protrusion-formation activity of MYO3A, then autophosphoregulation might be occurring at filopodial initiation sites on the inner surface of the plasma membrane. Such phosphorylation might not require MYO3A motor activity for the MYO3A kinase domain to bind and regulate the activity of other MYO3A molecules. Previous reports have shown that MYO3A kinase-alone truncations are capable of phosphorylating MYO3AΔK constructs in *in vitro* assays (Komaba et al., 2010; Quintero et al., 2010). To test the hypothesis that MYO3A phosphoregulation does not require motor activity to influence protrusion-initiation, we transfected COS7 cells with mchr-MYO3AΔK^WT^ in combination with GFP-MYO3A^FL^, or a kinase-domain-only truncation of MYO3A (GFP-MYO3A^KIN^). Unlike MYO3A constructs that contain an active motor domain (Quintero et al., 2010), GFP-MYO3A^KIN^ does not concentrate at filopodial tips (Figure 6). We then assayed the filopodia edge density of these cells as well as TCBR. Cotransfection of either GFP-MYO3A^FL^ (2.5±0.4, n=5, p<0.002, Tukey analysis) or GFP-MYO3A^KIN^ (3.1±0.1, n=5, p<0.02, Tukey analysis) with mchr-MYO3AΔK^WT^ resulted in significant decrease in TCBR compared to mchr-MYO3AΔK^WT^ alone (4.4±0.2, n=5, Figure 7A). Cotransfection of either GFP-MYO3A^FL^ (0.017±0.005 filopodia/μm, n=5, p<0.007, Tukey analysis) or GFP-MYO3A^KIN^ (0.021±0.003 filopodia/μm, n=5, p<0.02, Tukey analysis) with mchr-MYO3AΔK^WT^ also resulted in significant decrease in filopodia edge density compared to mchr-MYO3AΔK^WT^ (0.042±0.005 filopodia/μm, n=5, Figure 7B). This implies that intermolecular phosphorylation can occur in cells independently of motor function, and in other compartments in addition to the tips of actin protrusions. These data suggest that phosphoregulation of MYO3A could control the density of actin protrusion initiation events by regulating the amount of active MYO3A at filopodial initiation patches on the inner surface of the plasma membrane, in addition to regulating MYO3A activity at the tips of actin protrusions.

**Figure 6.**
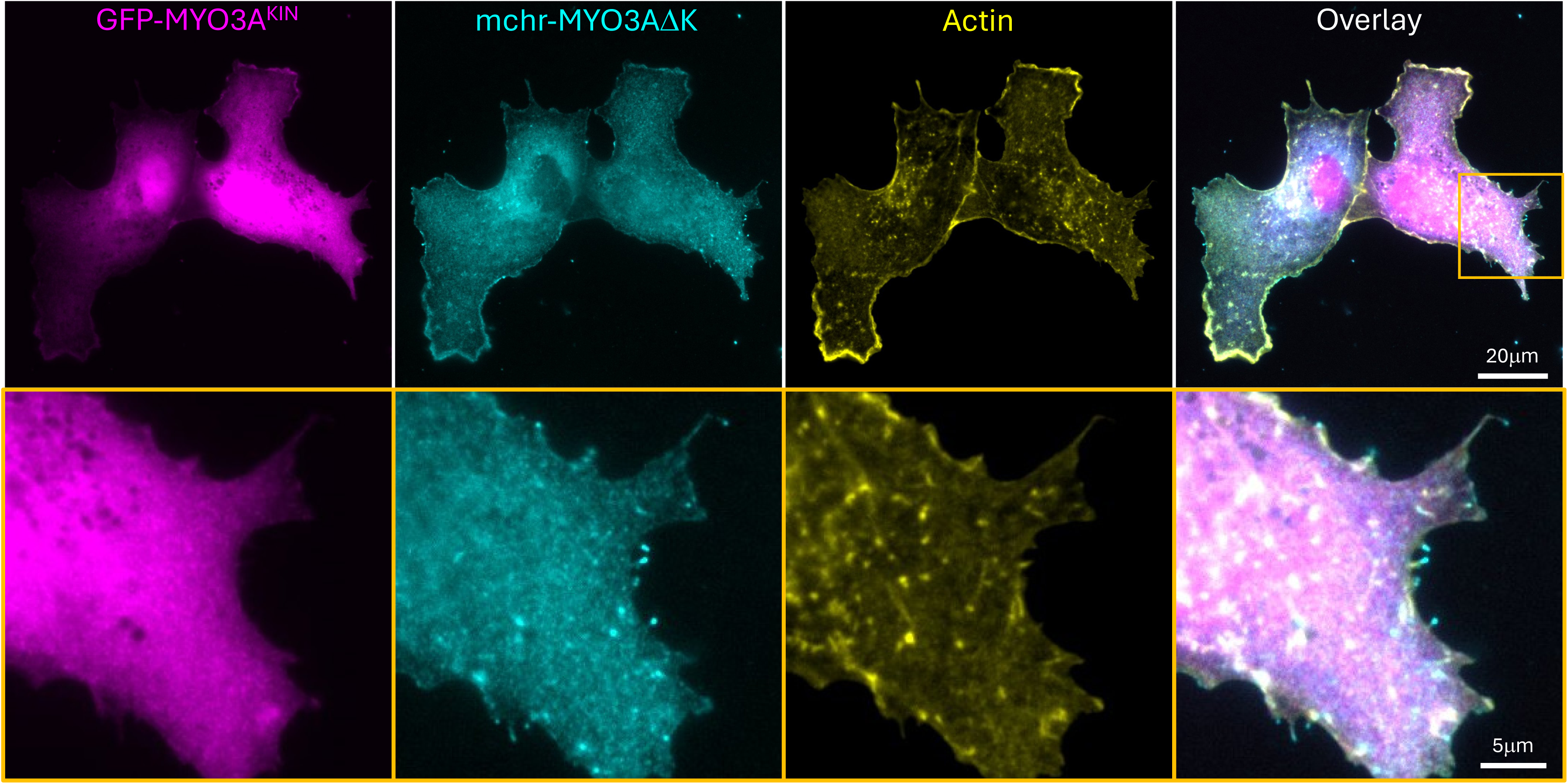
GFP-MYO3A^KIN^ does not localize to filopodial tips. **(A)** When mchr-MYO3AΔK (magenta) and GFP-MYO3A^KIN^ (cyan) are coexpressed in COS7 cells, mchr-MYO3ADK shows cytosolic localization and strong localization to filopodial tips. However, GFP-MYO3A^KIN^ does not concentrate at filopodial tips, remaining primarily cytosolic. The lower row if images are higher magnification representations of the region outlined in orange in the upper row of images. ALEXA647 phalloidin staining for F-actin is shown in yellow.

**Figure 7.**
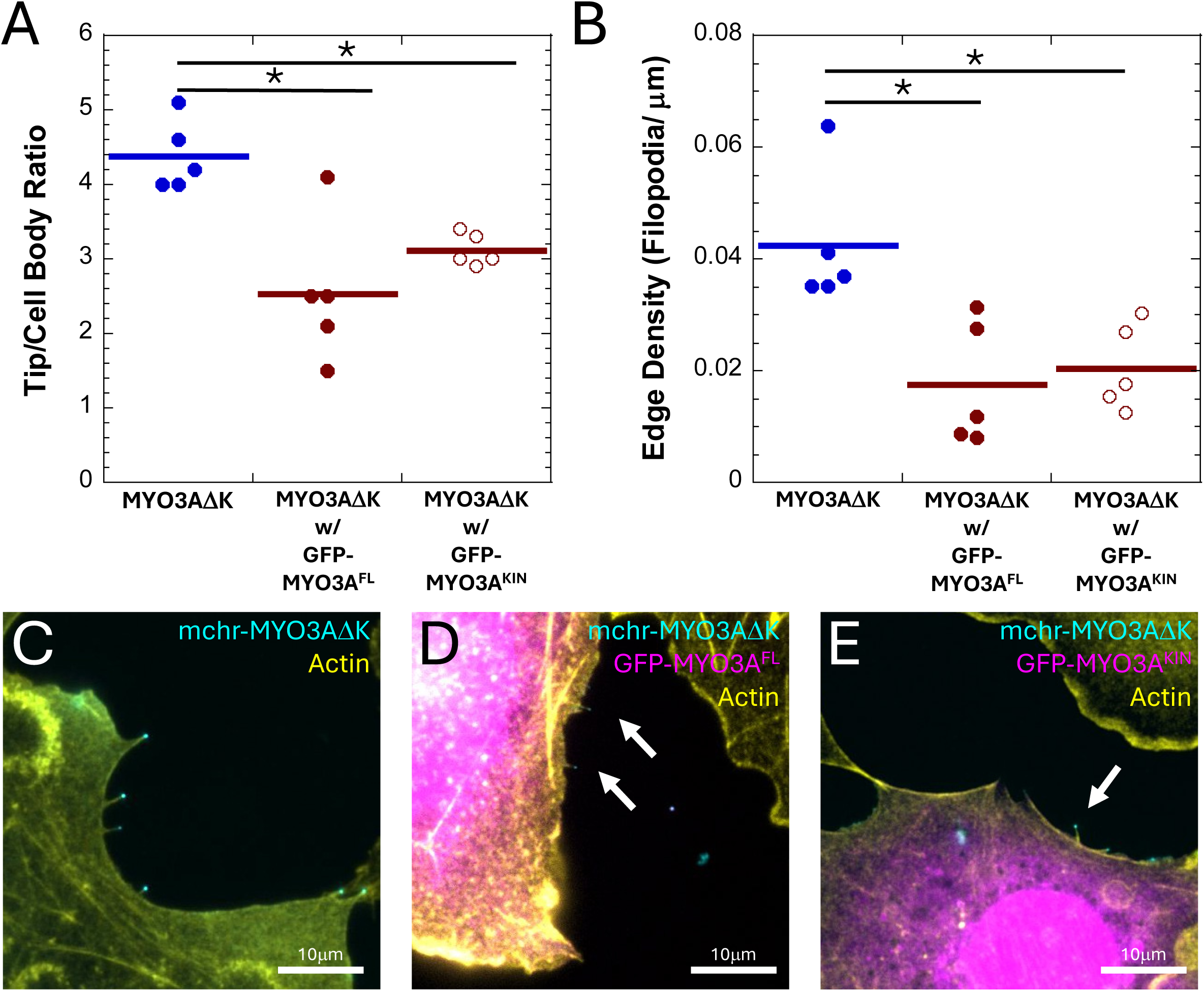
Coexpression of mchr-MYO3ΔK with GFP-MYO3A^KIN^ or GFP-MYO3A^WT^ results in decreased mchr-MYO3ΔK labeling at the tips and decreased filopodial edge density. **(A)** COS7 cells coexpressing mchr-MYO3ΔK with GFP-MYO3A^KIN^ have decreased tip localization and **(B)** decreased filopodial edge density when compared to mchr-MYO3ΔK^WT^ alone. COS7 cells coexpressing GFP-MYO3A^FL^ and mchr-MYO3AΔK also show a similar pattern of decreased tip localization and filopodial edge density. **C-E** are representative images of transfected cells. For all plots, markers represent biological replicates, bars represent the mean, and asterisks represent Tukey analysis with p<0.05.

To better understand the impact that phosphorylation at T908 and T919 might have on actin/MYO3A interactions, we threaded the human MYO3A sequence to a model structure for the actomyosin interface created by Lorenz and Holmes (Lorenz & Holmes, 2010) (Figure 8). Actomyosin interactions are partially dependent on electrostatic interactions between the two molecules, and this model predicted that both phosphorylation sites are likely positioned within the MYO3A/actin interface in close proximity to acidic patches containing aspartic acid residues D24 and D25 on actin. Phosphorylation would then introduce even more negative charge near these acidic residues, which in turn would decrease MYO3A’s affinity for actin, negatively influencing motor activity, and/or actin protrusion initiation. Previous studies on the motor kinetics of MYO3A have observed an overall decrease in the steady-state actin affinity of MYO3A (K_actin_) upon phosphorylation. Additionally, phosphorylation led to decreased maximal ATPase rate (k_cat_) and an increase in the actin concentration at which the ATPase rate was half-maximal (K_ATPase_), resulting in a decrease in steady-state myosin activity (Dosé et al., 2008; Komaba et al., 2010).

**Figure 8.**
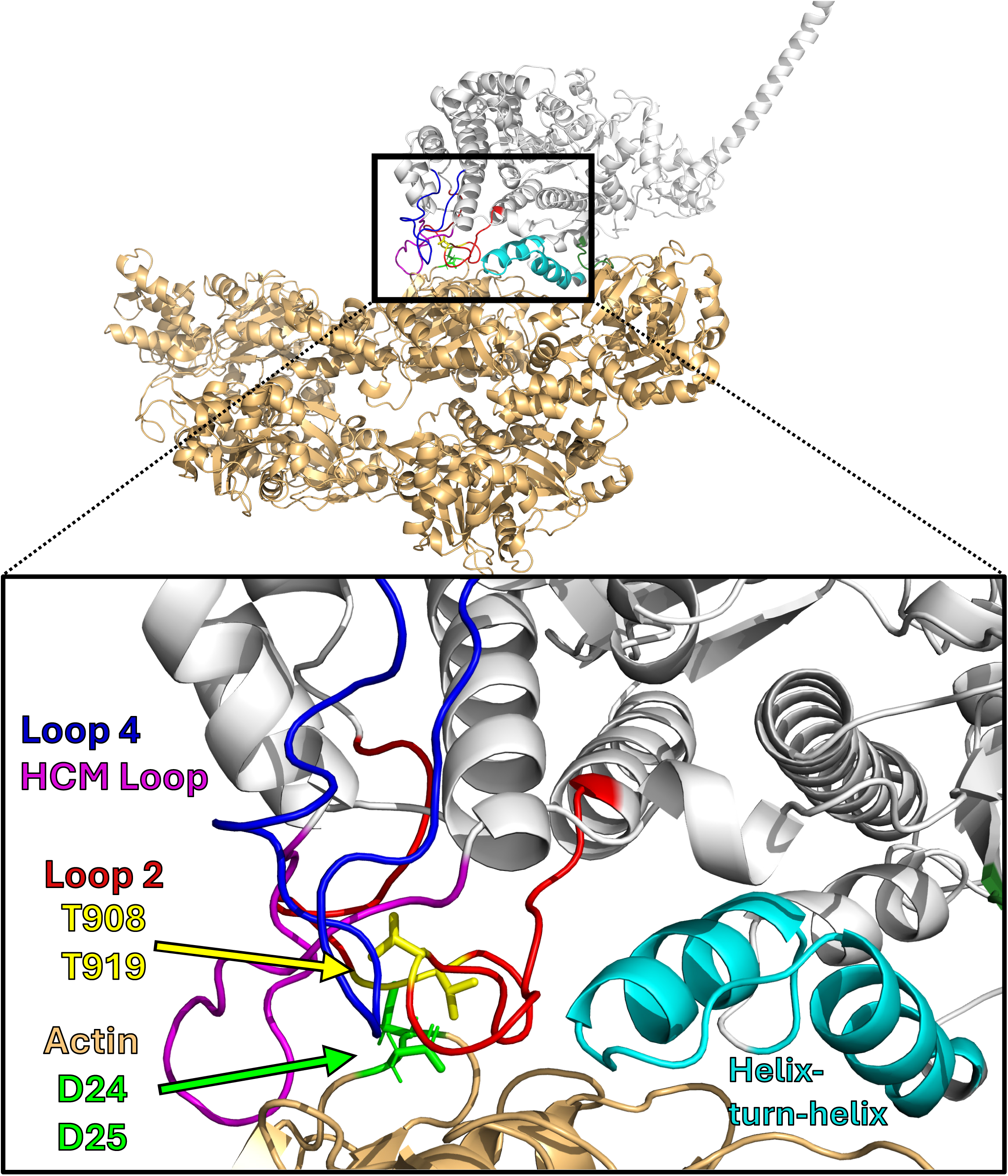
Structural predictions localize T908 and T919 residue near acidic residues on actin. The positions of residues T908 and T919 (highlighted in yellow) within the acto-MYO3A interface were predicted to be in proximity with acidic residues D24 and D25 on actin (highlighted in green). Phosphorylation at T908 or T919 will introduce negative charges that will lead to repulsion between the motor domain and actin. This model was generated by threading the human MYO3A sequence to a model of actin-bound MYH2 (Lorenz & Holmes, 2010) using Phyre2 (Powell et al., 2025).

Since intermolecular autophosphorylation can occur in the absence of MYO3A motor activity, MYO3A kinase activity could regulate actin-based protrusion initiation by MYO3A at nascent filopodial initiation sites along the plasma membrane. We propose a new model for MYO3A-mediated protrusion initiation where the amount of MYO3A present at protrusion initiation sites is determined by concentration-dependent MYO3A autophosphorylation (Figure 9). The higher the local MYO3A concentration, the more likely that the MYO3A in that cellular environment will phosphorylate each other. Active, unphosphorylated MYO3A collecting in patches at the membrane would induce actin protrusions, and as the local MYO3A concentration increased, MYO3A kinase activity would limit the concentration of active MYO3A available at the membrane to initiate a protrusion. Such a mechanism could regulate the size of protrusion initiation assemblies, influencing the density of actin-based protrusions. Such a phenotype has been observed in cells that generate filopodia (Quintero et al., 2010, 2013) and in cells that generate microvilli (Raval et al., 2016). Overexpression of human MYO3AΔK in hair cells resulted in elongated stereocilia that are characterized by higher amounts of MYO3A at the tips, and an overall “floppiness” that differs from the typical rigid stereocilia (Schneider et al., 2006). In this way, the presence of kinase activity tunes the motor domain’s activity, controlling actin-protrusion initiation activity.

**Figure 9.**
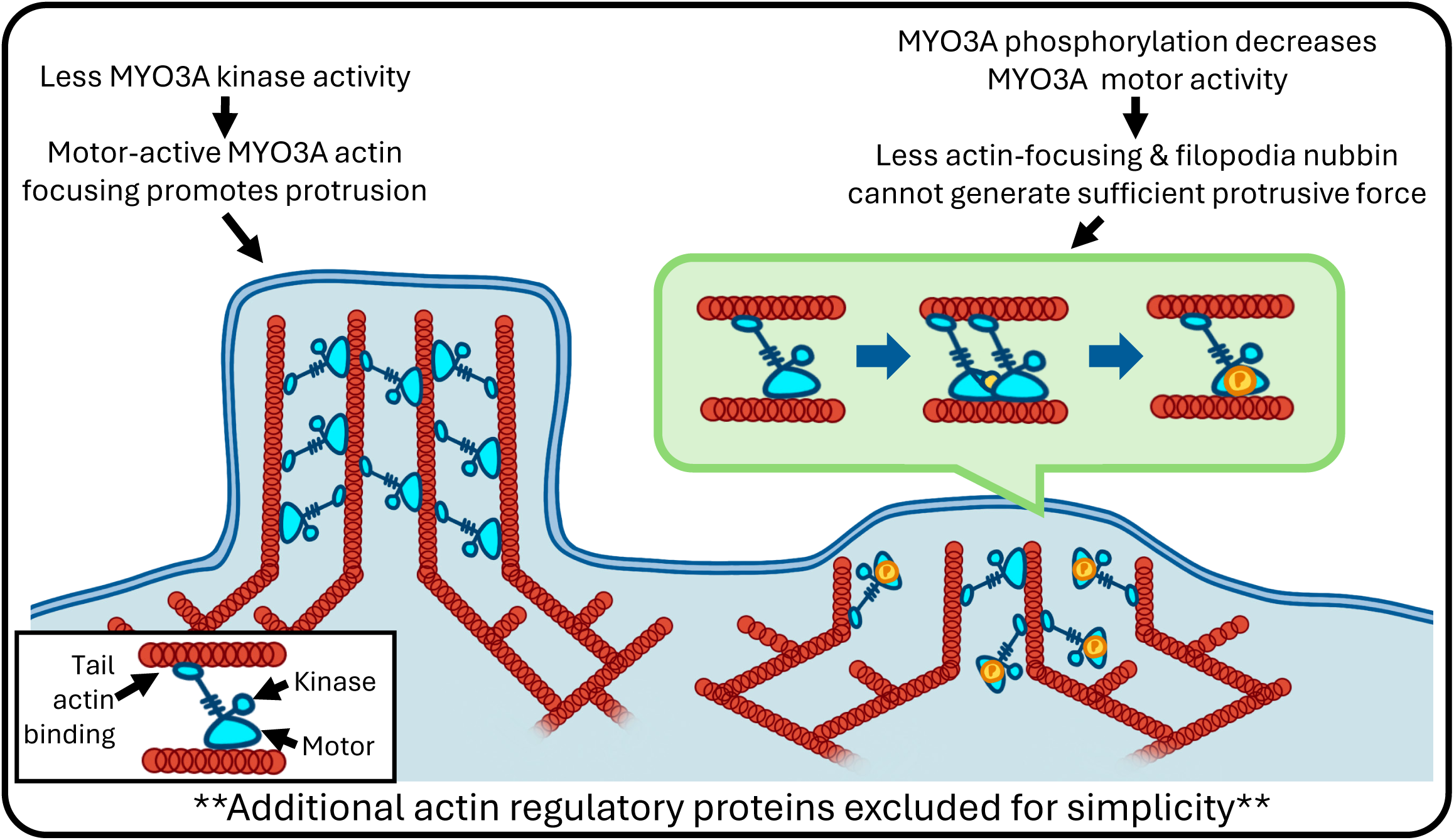
Model for actin-protrusion initiation mediated by phosphorylation of MYO3A. Decreased MYO3A kinase activity allows for more motor-active MYO3A actin-focusing activity to promote actin protrusions via actin elongation (left). Conversely, an increase in kinase activity results in phosphorylation of MYO3A, decreased motor activity and insufficient protrusive force generated by actin polymerization to support a nascent actin-protrusions (right).

## Conclusions and Future Directions

Our use of phosphomimicry has allowed us to investigate the biological effects of MYO3A phosphorylation at specific sites of the motor domain in a cell-based model system. Modeling human MYO3A to the Lorenz & Holmes model (Lorenz & Holmes, 2010) has predicted that upon phosphorylation at the T908 and T919 sites, there will be the presence of a negative where the motor domain would bind to negatively charged actin (Figure 8). Although we have not determined the molecular mechanism of phosphoregulation of the motor domain hypothesized to decrease in tip-localization in mchr-MYO3AΔK^T908D^ and mchr-MYO3AΔK^T919D^ constructs, the structural modeling and previous *in vitro* analyses suggest that a decrease in actin-activated myosin activity leads to a decrease in the ability of MYO3A to localize to and initiate actin-based protrusions. As MYO3A is a part of the myosin superfamily, it is capable of converting the energy from ATP hydrolysis into force and motion through a cyclic interaction with actin filaments. Although previous studies demonstrated that phosphorylation does alter the properties of the MYO3A motor *in vitro*, those studies looked at global motor domain phosphorylation and were not specifically addressing the changes to any particular phosphorylation site like phosphomimicry would. Future studies using MYO3A motor domain phosphomimics for *in vitro* analysis of steady-state and transient myosin kinetic behavior could lend further support to our hypothesis that T908 and T919 phosphorylation decrease MYO3A motor activity, and whether there is a difference in impact between the two phosphorylation sites.

Importantly, phosphorylation is a biological process that can happen whether or not a protein itself has a kinase or not. Given this, it is possible that MYO3A can be phosphorylated by kinases besides the MYO3A kinase domain through mechanisms beyond autophosphorylation. To test this, we could overexpress GFP-MYO3AΔK within COS7 cells, isolate the expressed protein via a pull-down assay with α-GFP antibodies (Yilmazer et al., 2023), and then run western blots for the immunoprecipitated samples using α-phosphothreonine antibodies to detect phosphorylation. Through this assay, we could identify if other kinases that might be components of additional regulatory pathways could regulate MYO3A activity. If other potential kinase activity was identified, then these preliminary data would support further investigation of other kinase signaling pathways in hair cells.

So far, we have only determined the individual effects of phosphorylation at T908 and T919 regions within the motor. As both sites could be phosphorylated simultaneously, accumulation of phosphorylations could have additive effects not observed in the single phosphomimic constructs. Through generation of double phosphomimic construct, we could determine whether double phosphorylation results in stronger inactivation of MYO3A filopodial initiation activity. Additionally, the generation of individual phosphonulls (T908A or T919A) and double phosphonulls (T908A in combination with T919A) that are resistant to phosphorylation could also reveal whether these two sites are the phosphorylation sites primarily responsible for regulating MYO3A motor function.

Changing an amino acid sequence can have drastic effects on protein structure and by extension, its function. To determine whether the observed difference between mchr-MYO3AΔK and the phosphomimics mchr-MYO3AΔK^T908D^ and mchr-MYO3AΔK^T919D^ are due to the imitation of phosphorylation and not solely due to a change R-group properties, it may be beneficial to generate T908S and T919S point mutations. Serine is similar in chemical nature to threonine. Although serine can also be phosphorylated, it is likely that the MYO3A kinase domain would not phosphorylate serine substitutions readily as its R-group is smaller. Using such constructs in our cell-based assays or *in vitro* myosin activity assays could provide insights as to the importance of phosphothreonines and potentially the selectivity of the MYO3A kinase domain.

### Further exploring the biological relevance of the other domains of MYO3A

MYO3A has been hypothesized to play a role in the initiation of stereocilia, possibly through its transportation of ESPN1 (Dosé et al., 2007). Currently, our metric of filopodial edge density cannot differentiate between increases in filopodia initiation, increases in filopodia stabilization, or decreases in rates of filopodia loss mediated by MYO3A. Live-cell imaging approaches using COS7 cells expressing mchr-MYO3AΔK^WT^ would reveal whether these constructs influenced how often filopodia formed, how fast they extended, and how long they persisted. Cotransfection with GFP-MYO3A^FL^ or GFP-MYO3A^KIN^ would then allow us to determine the effects phosphorylation on those kinetic measures of filopodia behavior, and using mchr-MYO3AΔK^T908D^ and mchr-MYO3AΔK^T919D^ constructs in live-cell assays could reveal the impact of phosphorylation at particular sites on filopodia behavior.

The motor domain of MYO3A is not the only region within the protein that has potential phosphorylation sites. Previous studies by Quintero et al., have shown phosphorylation also occurs at T184 of the MYO3A kinase domain. This site is part of the kinase domain “activation loop,” (Quintero et al., 2013) and T184 phosphorylation enhances the phosphorylation ability of the kinase domain. Phosphoproteomic data in their report as well as other reports have also noted additional phosphorylation sites within the tail domain of MYO3A that have yet to be studied. Characterization of these other potential phosphoregulatory sites could alter the affinity of ESPN1 and/or MORN4 binding to MYO3A, impacting the biological function of the entire assembly through a mechanism unrelated to motor function.

A previous model on the phosphoregulation of MYO3A proposed that MYO3A undergoes <u>c</u>oncentration-<u>d</u>ependent <u>r</u>egulation by <u>a</u>uto<u>p</u>hosphorylation (CD-RAP) (Quintero et al., 2010). In this model, accumulation of MYO3A at the tips of actin protrusions increases the likelihood of intermolecular autophosphorylation resulting in decreased motor activity. This reduction in motor activity then leads to regulation of MYO3A activity in that particular compartment (Quintero et al., 2010). However, our studies indicate that phosphoregulation of MYO3A may occur wherever MYO3A accumulates, whether that is in the tip of established filopodia or along the cell membrane where actin-based protrusions are forming. The transport of the kinase domain to the tips via the motor is not necessary for the regulation of MYO3A, as we see a decrease in filopodial edge density and tip localization for mchr-MYO3AΔK^WT^ when coexpressed with GFP-MYO3A^KIN^. Additionally, phosphoregulation may alter the behavior of other MYO3A properties besides motor activity.

As such we proposed a new model, where in addition to CD-RAP at the tips, the regulation of MYO3A can happen within the cell body through the kinase activity resulting in phosphorylation of the motor domain (Figure 9). This sort of kinase activity could control the ability of tip-complex assemblies containing MYO3A, ESPN1, MORN4, actin, or other necessary components to generate the force needed to promote polymerization-based initiation and elongation of actin protrusions.

## VII. Acknowledgments

Thank you to the University of Richmond School of Arts & Science for funding and support, as without this, I do not think I would have been able to pursue this passion for science. Thank you to all the Q-Lab members, Chase Cristella, Anna Lozano, Joanna Mas, and Lillie Wendt whom I’ve had the honor of being a Q-labby with. Inside and outside of the lab, you guys have shown me how wonderful and whimsical life can be and I will truly miss seeing you guys around. Special shoutout to Lillie specifically for her immense artistic abilities and drawing Figure 9. I would also like to thank Dr. Stacey Criswell for her never-ending patience and wonderful mentoring while teaching me how to use basically every microscope in Gottwald. You have allowed me to see the world in an entirely different way, and for that I am forever grateful. Finally, I would like to thank Dr. Omar Quintero-Carmona, for without his influence and unconditional support, I don’t think I would have been able to find the same joy and fulfillment in science. In my time at University of Richmond, no words brought me greater joy than you asking me, “do you want to see some cells?”, and I will miss that dearly as I try to make you proud on this path you have helped set me on.

